# Classification of clear cell renal cell carcinoma based on *PKM* alternative splicing

**DOI:** 10.1101/823336

**Authors:** Xiangyu Li, Beste Turanli, Kajetan Juszczak, Woonghee Kim, Muhammad Arif, Yusuke Sato, Seishi Ogawa, Hasan Turkez, Jens Nielsen, Jan Boren, Mathias Uhlen, Cheng Zhang, Adil Mardinoglu

## Abstract

Clear cell renal cell carcinoma (ccRCC) accounts for 70–80% of kidney cancer diagnoses and displays high molecular and histologic heterogeneity. Hence, it is necessary to reveal the underlying molecular mechanisms involved in progression of ccRCC to better stratify the patients and design effective treatment strategies. Here, we analyzed the survival outcome of ccRCC patients as a consequence of the differential expression of four transcript isoforms of the pyruvate kinase muscle type (*PKM*). We first extracted a classification biomarker consisting of eight gene pairs whose within-sample relative expression orderings (REOs) could be used to robustly classify the patients into two groups with distinct molecular characteristics and survival outcomes. Next, we validated our findings in a validation cohort and an independent Japanese ccRCC cohort. We finally performed drug repositioning analysis based on transcriptomic expression profiles of drug-perturbed cancer cell lines and proposed that paracetamol, nizatidine, dimethadione and conessine can be repurposed to treat the patients in one of the subtype of ccRCC whereas chenodeoxycholic acid, fenoterol and hexylcaine can be repurposed to treat the patients in the other subtype.

## Introduction

Clear cell renal cell carcinoma (ccRCC) is the most common subtype among renal cancer (Motzer et al., 2017) and ccRCC shows high inter individual heterogeneity(Ricketts et al., 2018). Thus, it is difficult to predict survival outcomes of patients in clinical practice and design effective therapeutic strategies. Previous studies have already proposed strategies for stratification of ccRCC patients into subgroups based on different genetic and/or transcriptomic characteristics and prognoses of the patients (Brannon et al., 2010; Cancer Genome Atlas Research, 2013; Kosari et al., 2005; Takahashi et al., 2001). However, these studies failed to identify a clinically practical biomarker for classification of the patients at the personalized level or recommend personalized chemotherapy regimens for these patients.

In a recent study, we have found that pyruvate kinase muscle type (*PKM*), an enzyme that is involved in the final step of glycolysis and catalyzes the formation of ATP from ADP as phosphoenolpyruvate undergoes dephosphorylation to pyruvate, plays a very important role in controlling tumor metabolism in ccRCC (Li et al., 2019b). We have also observed that the expression level of four protein-coding transcripts of *PKM*, including ENST00000335181, encoding PKM2 which is the most studied isoform of PKM, as well as ENST00000561609, ENST00000389093 and ENST00000568883 are highly associated with patients’ prognoses. Among them, high expression of ENST00000335181 and ENST00000561609 indicate a favorable survival while high expression of ENST00000389093 and ENST00000568883 indicate an unfavorable survival. Moreover, a number of conserved biological functions associated with the progression of ccRCC were oppositely dysregulated by these transcripts. Here, we hypothesized that different molecular subtypes among ccRCC patients may be characterized by the different expression patterns induced by these four prognostic transcripts and biomarkers that may be used in clinical practice can be identified.

Previous studies have proposed transcriptomics-based biomarkers for classification of tumors based on the quantitative measurement of one or multiple signature genes (Fujita et al., 2012; Jones et al., 2005; Klatte et al., 2009; Kosari et al., 2005; Zhao et al., 2006). However, this kind of transcriptional signatures are rarely used in clinical practice due to technological and translational barriers (Winslow et al., 2012). Besides problems in tissue sampling and quality control, an important factor is experimental batch effect which brings high variation of gene expression induced by the different laboratory conditions and personnel (Guan et al., 2018). To solve these problems, the use of biomarkers based on the within-sample relative expression orderings (REOs) of gene pairs has been proposed (Guo et al., 2018; Qi et al., 2016a; Qi et al., 2016b), which is robust against batch effects, invariant to monotone data normalization (Eddy et al., 2010; Wang et al., 2013) and poor sample preparation (Chen et al., 2017b; Cheng et al., 2017; Liu et al., 2017).

In this study, we used the genes dysregulated by the prognostic transcripts of *PKM* to extract classification biomarker instead of using themselves. Since different transcripts of *PKM* share similar sequence, it may be difficult to design distinct primers to detect their relative abundance when using real-time PCR. Thus, gene pair biomarker is more feasible and practical in clinical diagnosis. We applied REOs-based method to identify classification biomarker for ccRCC by extracting the expression profiles of genes which were consistently negatively dysregulated by the four favorable and unfavorable prognostic transcripts of *PKM*. We developed a REOs-based biomarker using the global gene expression profiling of ccRCC in The Cancer Genome Atlas (TCGA) database and stratified the patients into two subtypes exhibiting different transcriptomic expression patterns and different patient prognosis. We also validated our findings in TCGA database as well as in another independent Japanese cohort. We finally proposed several candidate drugs that can be used in treatment of each subtype based on transcriptomic expression profiles of drug-perturbed cancer cell lines from Connectivity Map 2.0 (CMap2).

## Result

### Identification of signature gene set associated with four prognostic transcripts of *PKM*

In a recent study, we have found that there are molecular subtypes that could be characterized by the expression of the four prognostic PKM transcripts (Li et al., 2019b). In order to develop a REOs-based biomarker, we identified a signature gene set associated with these four transcripts based on the gene expression profiles of TCGA ccRCC samples. We performed differential expression analysis between the tumor samples from patients with high (top 25%) and low (bottom 25%) expression of each favorable transcript, and identified 2010 consistently significantly (FDR < 1.0e-5) differentially expressed genes (DEGs) for the two favorable transcripts (Figure 1). Similarly, we identified 5469 DEGs consistently significantly (FDR < 1.0e-5) DEGs for the two unfavorable transcripts. We found that the two sets of DEGs has a significant overlap (n=1135; hypergeometric distribution test, p<1.11e-16). We also observed that the concordance score of these overlapped genes is 100%, which means the up-regulated genes associated with high expression of favorable transcripts within these 1135 genes are all down-regulated when the unfavorable transcripts exhibit high expression; and vice versa.

**Figure 1.**
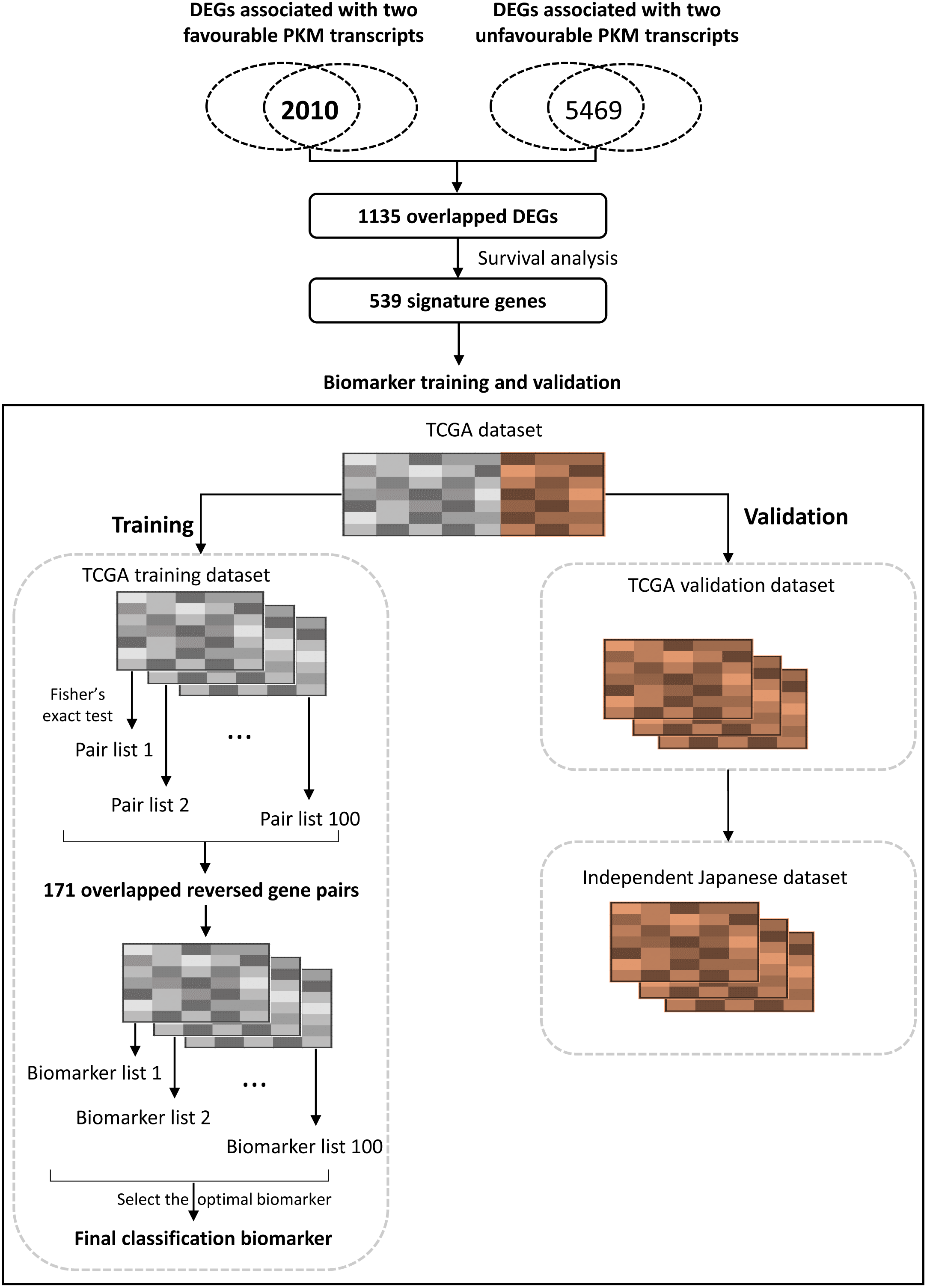
The flowchart for developing and validating the ccRCC classification biomarker. In brief, we extract the 1135 overlapped DEGs associated with favorable and unfavorable transcripts, and select 539 prognostic signature genes from them. Next, we screen gene pair biomarker using randomly generated training dataset. Lastly, we validate the performance of the biomarker in all randomly generated validation dataset and an independent Japanese ccRCC dataset.

We followed-up survival information from the corresponding patients and found 539 of the 1135 genes (of which 305 and 234 are favorable and unfavorable, respectively) are significantly (univariate Cox model, FDR < 0.01) associated with patients’ overall survival (OS). To identify the associated biological functions with these 539 genes, we performed GO term enrichment analysis and observed that these genes are significantly enriched in RNA splicing, RNA catabolic process and nuclear transport pathways (FDR<0.05; Table S1). Therefore, we concluded that these 539 genes may be used as the core signature genes that are associated with the differential alternative splicing of *PKM* among ccRCC patients and may be used for classification of tumor samples.

We calculated the co-expression coefficients between the expression of the 539 signature genes and found two major clusters in which all favorable genes are positively co-expressed while all unfavorable genes are negatively co-expressed in the opposite cluster using the hierarchical clustering (Figure 2A). Based on the expression profiles of these 539 signature genes, we employed consensus clustering to classify TCGA ccRCC samples into distinct stable groups through repeated subsampling and clustering (Wilkerson and Hayes, 2010). As shown in Figure 2B, we determined an optimum number of two clusters, cluster 1 and 2, based on the lowest proportion of ambiguous clustering (Senbabaoglu et al., 2014). Using survival analysis, we observed the patients whose tumor samples classified in cluster 1 (N = 231) had significantly shorter OS than those classified in cluster 2 (N = 297) with statistical significance (log-rank test, P=6.73e-07; Figure 2C). The result demonstrated that there are two different molecular subtypes in ccRCC with significantly different survival outcomes which are strongly associated with the function of the two favorable and two unfavorable transcripts.

**Figure 2.**
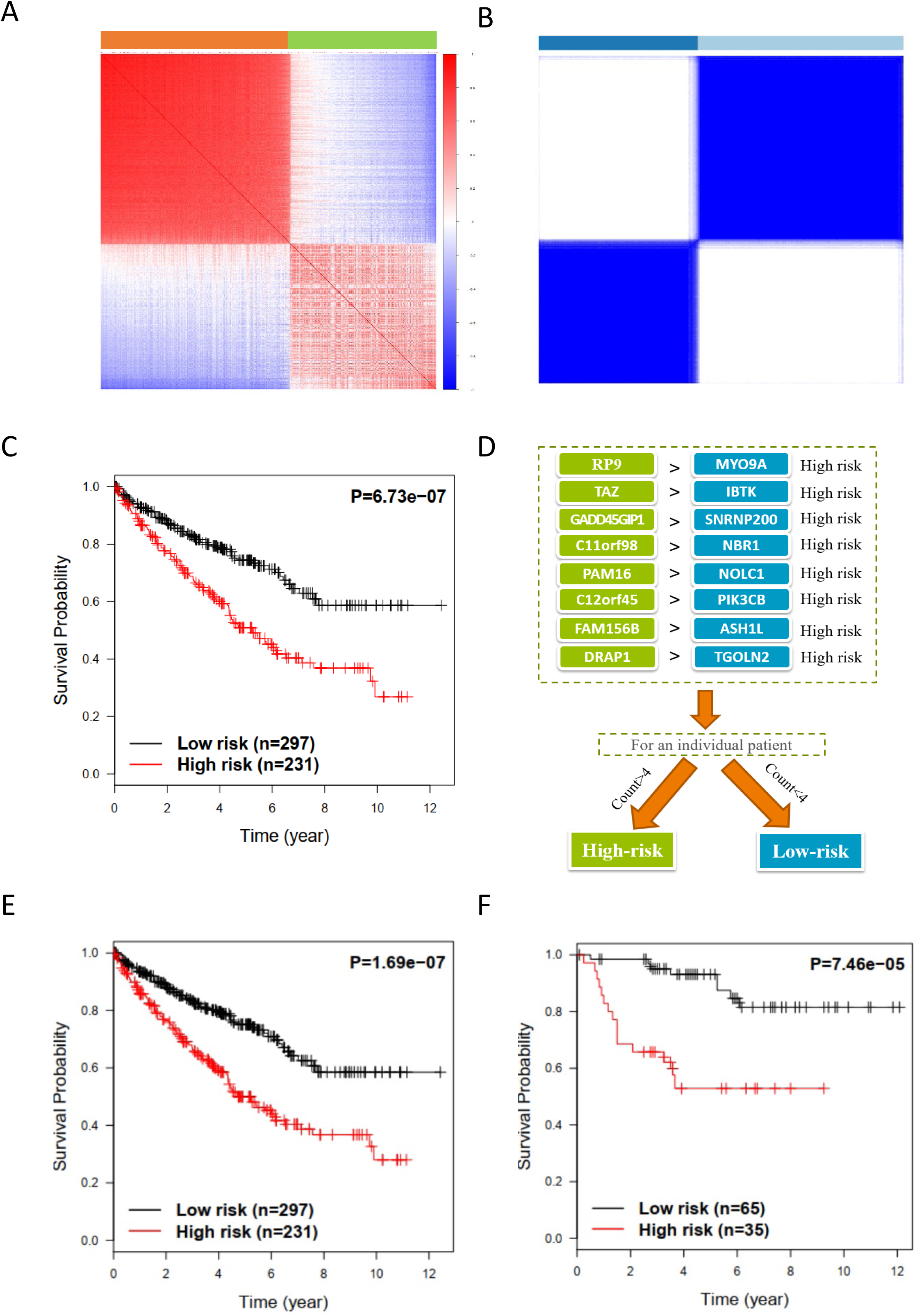
Molecular classification and prognostic prediction of patients by classification biomarker. (A) Hierarchical clustering of 539 signature genes based on the correlation between genes. The spearman correlation coefficients between genes were used for clustering. (B) Consensus clustering for TCGA ccRCC patients based on the expression values of the 539 signature genes. (C) Kaplan-Meier plot of OS of two clusters identified by consensus clustering in TCGA ccRCC cohort. (D) The composition of classification biomarker and voting rule. (E) Kaplan-Meier plot of OS of high- and low-risk identified by classification biomarker in TCGA ccRCC cohort. (F) Kaplan-Meier plot of OS of high- and low-risk identified by classification biomarker in Japanese ccRCC cohort.

### Development of the REOs-based classification biomarker

To identify a biomarker that can be used in the clinical practice, we next focused on development of a REOs-based classification biomarker based on the gene expression profiles of the 539 signature genes. In brief, REOs-based biomarkers employs gene pairs with consistently reversed expression orders between the two molecular groups as indicators, and screens for a minimum combination of these gene pairs that serves as risk indicators for classification. In order to obtain a robust biomarker, we generated 100 training and 100 validation datasets by randomly selecting from TCGA ccRCC cohort and randomly separated the samples into two respective groups with 70% and 30% samples. We identified 171 gene pairs that exhibited consistent reverse gene pairs in all training datasets. We next generated 17100 reverse gene pair combinations with a forward selection procedure and selected a final REOs-based biomarker consisted of eight reverse gene pairs with an optimal mean F-score of 0.9725 in all training datasets. The full screening process are shown in Figure 1 and detailed in Method section.

Within this eight gene pairs, if more than four gene pairs exhibited reversal REOs in a sample, this sample would be classified into the high-risk group; otherwise, this sample would be classified into the low-risk group (Figure 2D). We tested these gene pairs in the 100 validation datasets, and found that these gene pairs also showed a good classification accuracy with a mean F-score 0.9742. We also tested these gene pairs using the complete TCGA cohort, and this biomarker classified 231 samples into high-risk group and 297 samples into low-risk group. Notably, these two groups showed significantly different OS (Figure 2E; log rank test, P=1.69e-07).

### Validation of the REOs-based classification biomarker

To validate if these gene pairs can be used as a biomarker for classification of ccRCC samples, we tested these gene pairs in 100 ccRCC samples obtained from an independent Japanese cohort. The biomarker classified 35 samples as high-risk group and 65 samples as low-risk group, and these two groups showed significantly different OS (Figure 2F, P=7.46e-05). We next investigated whether the high and low-risk groups identified in both TCGA and Japanese cohorts exhibited similar biological differences. We extracted the top 20% most significant DEGs (n=2694) between high and low-risk groups in both the TCGA and Japanese cohorts, and observed a significant overlap between them (*n* =1463; hypergeometric distribution test, *p* < 1.11e-16) with a concordance score 100%. In addition, we identified 66 and 80 GO terms that are significantly enriched with upregulated genes (FDR<1.0e-05) in the high-risk group of the TCGA and Japanese cohorts, respectively, and found that 55 of them are common in both cohorts (Figure 3). Specifically, the high-risk group was characterized by upregulated genes involved in ATP synthesis, mitochondrial respiratory process, oxidative phosphorylation, ribonucleotide and purine nucleotide metabolic process, RNA catabolic process, protein targeting to ER and membrane pathways. And the low-risk group was characterized by upregulated genes involved in histone modification and covalent chromatin modification pathways. The results suggested the molecular subtypes identified by our analysis also have consistent biological differences. Moreover, these 55 GO terms included all 27 GO terms that we recently reported to be associated with the four prognostic transcripts in pan cancer analysis (Li et al., 2019b). We also identified three GO terms that are significantly enriched with downregulated genes in the high-risk group for both cohorts, and two of them, which are the histone modification and covalent chromatin modification pathways, are common in both groups. These results further indicated that the molecular subtypes stratified by the gene pairs are functionally related to the four prognostic transcripts of *PKM.*

**Figure 3.**
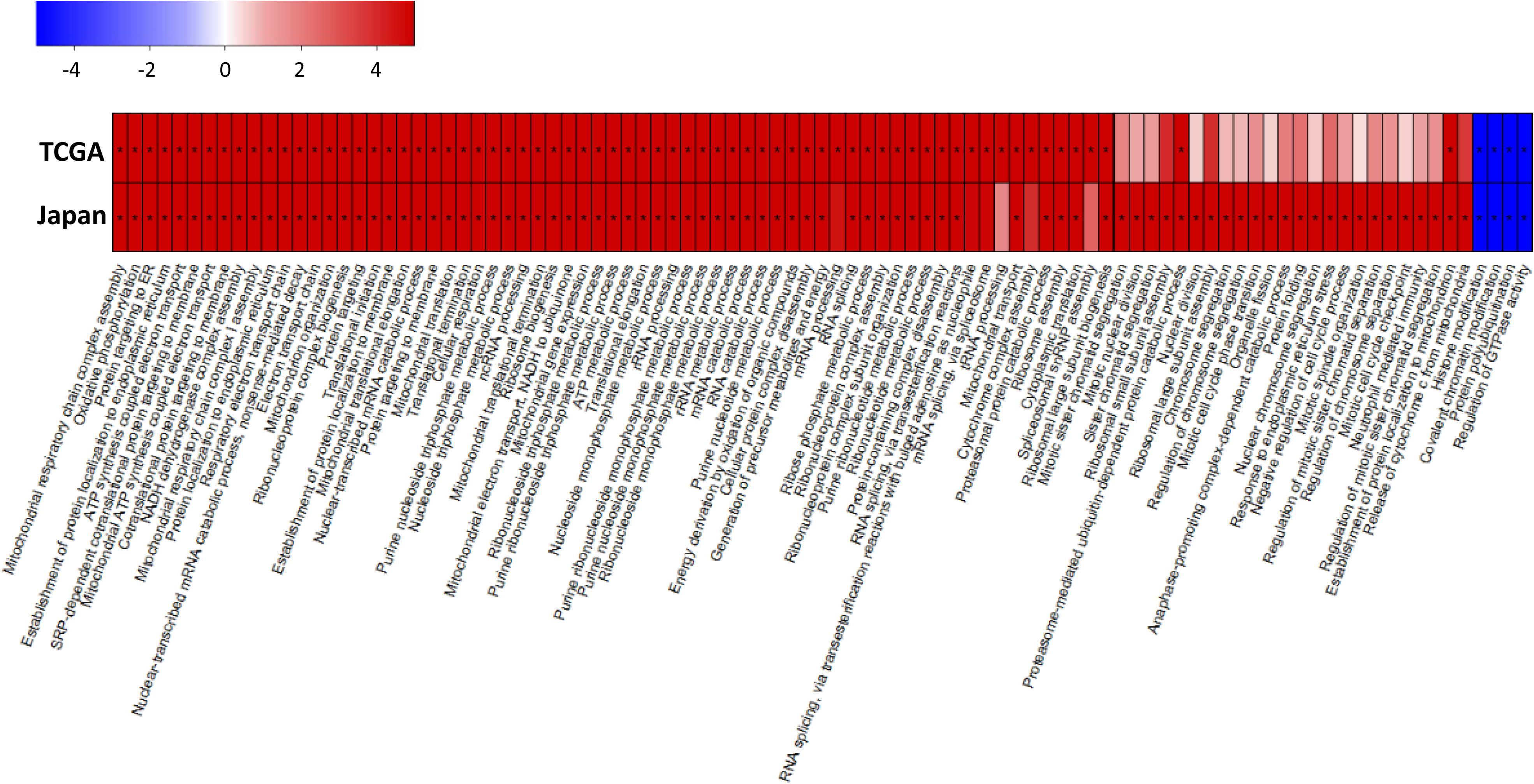
The dysregulated biological functions in high- and low-risk ccRCC groups and. Heat map of the p values (on the negative log 10 scale) for the enriched GO terms in TCGA and Japanese KIRC cohort. Red color denotes the GO terms enriched with up-regulated genes. Blue color denotes the GO terms enriched with down-regulated genes. * FDR<1.0e-05.

Further, we compared our REOs-based classification with previously reported TCGA (m1 to m4) and ccA/ccB classification schemes (Brannon et al., 2010; Cancer Genome Atlas Research, 2013) (Figure 4). In TCGA cohort, approximately 96% of TCGA m1 tumors were involved in our low-risk group, and m1 group was also reported with the best prognoses in TCGA classification scheme. In addition, 73% of TCGA tumors in both m2 and m4 subtypes were involved in our high-risk groups, and they were also shown to be with the poorest prognoses in TCGA classification. These results demonstrated that the high and low-risk groups classified by our biomarker are reinforced by the previously observed survival outcomes m1, m2 and m4. Notably, the high and low-risk groups respectively accounted for 42% and 58% of tumors previously reported as unclassified (m3) in the TCGA classification scheme. In Japanese cohort, 71% of ccA and 80% of ccB were observed in the low and high-risk groups, respectively. We found that the favorable survival for ccA cases again reinforced the low-risk group classification based on our gene pairs used as a biomarker (Figure 4).

**Figure 4.**
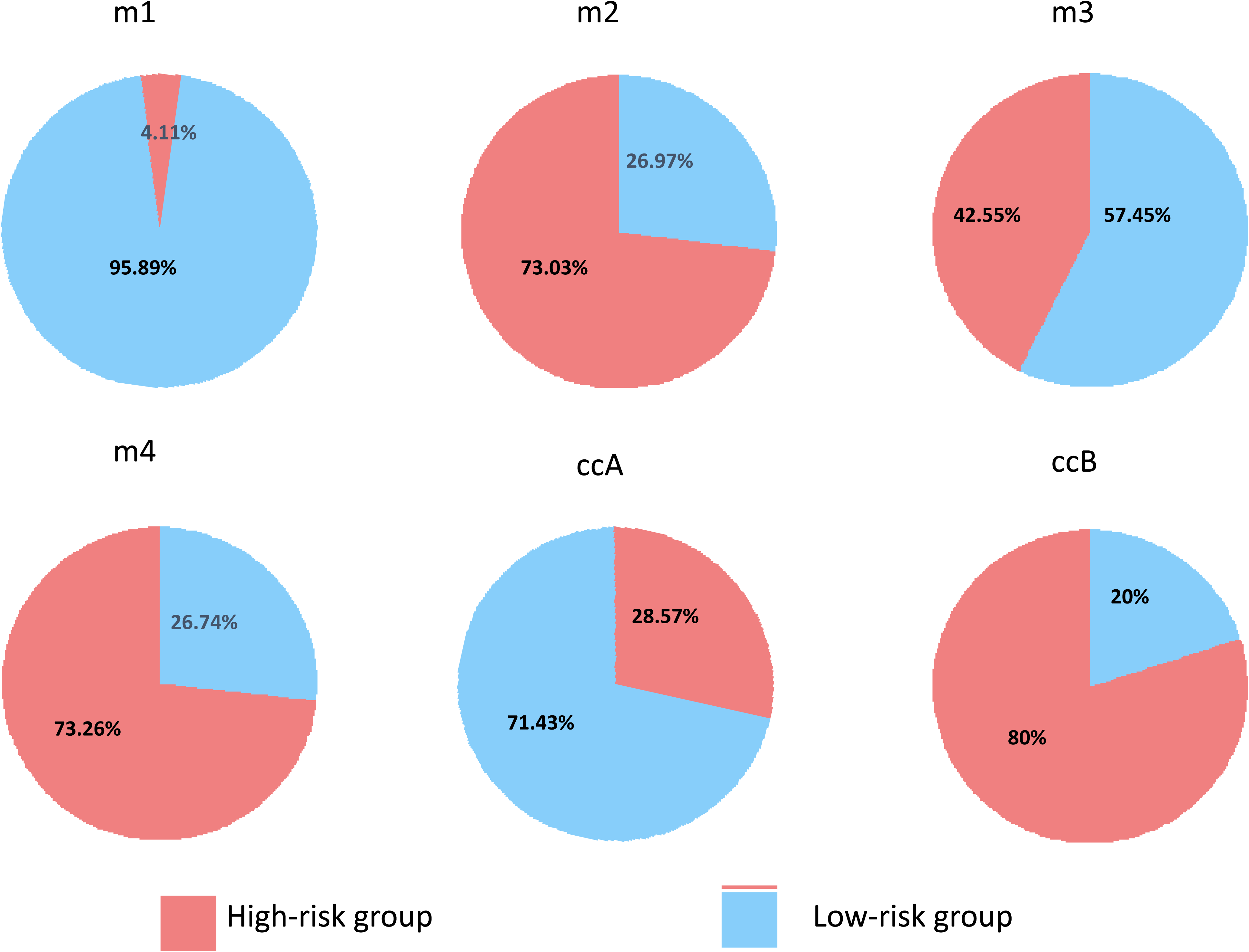
Pie charts showing the intersection of the different classification schemes for ccRCC. ‘m1’, ‘m2’, ‘m3’ and ‘m4’ indicate the molecular subtypes proposed by TCGA, and ‘ccA’ and ‘ccB’ are molecular subtypes reported by another previous study.

### Drug discovery with reversed expression effect

In addition to classification of the tumors, we also performed drug repositioning analysis to identify drug candidates that can be used in treatment of each subtype. We assumed that if a drug could reverse the dysregulated gene expression pattern from a tumor subtype to normal pattern, it could be potentially useful for treating the specific tumor subtype. We used a method developed in our previous study for drug repurposing (Turanli et al., 2019a; Turanli et al., 2018; Turanli et al., 2019b) and found several drugs that could be used for treatment of the high and low-risk groups. We found that four different drugs including paracetamol, nizatidine, dimethadione and conessine could be used to reverse the gene expression in samples from high-risk group, since over 80% drug-perturbed genes were mapped to the DEGs between these samples and normal samples. Interestingly, it has been reported that paracetamol, an analgesic and antipyretic drug, inhibits the cell proliferation and induces cell apoptosis in pancreatic cancer (Malsy et al., 2017), ovarian cancer and lung cancer cells (Lian et al., 2018). Nizatidine, a histamine H_2_ receptor antagonist, was also recommended to be added into the combination therapy for cancer treatment (Barton-Burke, 1996; Ben-Sasson, 2007; Feitelberg et al., 2013). Therefore, the anti-cancer effects of two of the proposed drugs have been validated in previous studies.

Similarly, we found that three different drugs including chenodeoxycholic acid, fenoterol and hexylcaine, could be used to reverse the gene expression in samples of low-risk group towards normal samples. It has been reported that chenodeoxycholic acid, a bile acid, shows anti-proliferative activity in human cancer cells (Faustino et al., 2016). Fenoterol, a β adrenoreceptor agonist, has been shown to inhibit proliferation of glioblastomas and astrocytomas cells (Bernier et al., 2013; Toll et al., 2011). Hexylcaine, a short-acting local anesthetic, has also been used to treat cancer (Gleich, 2000). In this context, these three drugs may also be potentially used for treatment of the subtype of ccRCC patients.

Moreover, we identified the gene targets for each of the drugs using DrugBank database (Wishart et al., 2006). *HRH2*, encoding the histamine H2 receptor, has been reported as the gene target of nizatidine (Meredith et al., 1985). We observed that it is significantly up-regulated in the samples of high-risk group compared to normal samples. It has been demonstrated that in vitro and in vivo histamine-induced tumor cell proliferation can be blocked by H2 antagonists (Deva and Jameson, 2012; Natori et al., 2005; Tomita et al., 2003). Thus, nizatidine may be used as a promising drug for the patients classified in the high-risk group. On the other hand, *GPBAR1*, encoding an enzyme of the G protein-coupled receptor superfamily, has been reported as a target of chenodeoxycholic acid. It has been shown that *GPBAR1* antagonizes kidney cancer cell proliferation and migration (Su et al., 2017). Based on our analysis, we have observed that *GPBAR1* is significantly downregulated in the low-risk group compared to normal samples. Thus, chenodeoxycholic acid, as an activator of *GPBAR1*, may be used as a promising drug for treating the patients classified in the low-risk group.

## Discussion

*PKM* is one of the important regulators of Warburg effect in different human cancers (Dayton et al., 2016). Our recent study showed that four different transcripts of PKM mediated opposite survival outcomes for ccRCC patients. In this study, we identified the core signature genes which were consistently dysregulated by these four prognostic transcripts of *PKM*. Using these signature genes, we identified eight gene pairs whose within-samples REOs could be used to classify patients into two groups with significantly different OS. REOs-based biomarkers take the advantages of the robustness of the intra sample gene expression pattern, and it is relatively insensitive to both experimental and bioinformatics variations compared to conventional biomarkers based on absolute quantification. Although RNA sequencing data was used for the biomarker classification in this study, much cheaper technics could be used once the biomarker is used in clinical practice. For instance, we could use real-time PCR, which is much cheaper compared to the sequencing approach, to determine the relative abundance of the genes involved in these 8 gene pairs to classify a ccRCC tumor sample since we only need to detect their REOs. This could greatly facilitate the use of REOs-based biomarker in clinical practice.

The genes involved in our classification of ccRCC tumor samples also showed closed relationship with tumor development. For instance, RP9, one of genes involved in the REOs gene pairs, plays an important role in pre-mRNA splicing and could interact with well-known oncogene PIM-1(Maita et al., 2000). TAZ, another gene involved in the REOs gene pairs, encodes tafazzin whose overexpression promotes tumorigenicity in many cancers and its inhibition also induces tumor cell apoptosis (Chen et al., 2017a; Li et al., 2019a; Pathak et al., 2014). Another example is NOLC1 which functions as a chaperone for shuttling between the nucleolus and cytoplasm (Meier and Blobel, 1992). It has been reported that enhancement of NOLC1 promotes cell senescence and represses hepatocellular carcinoma cell proliferation by disturbing the organization of nucleolus (Yuan et al., 2017).

In conclusion, we identified two molecular subtypes of ccRCC patients with high and low-risk of mortality, and developed a REOs-based classification biomarker which could be used to identify which subtype of the patients belong to in a personalized manner. In addition, we also suggested specific treatment strategies for each subtype based on their global gene expression patterns. Therefore, it is worthwhile to further explore the potential clinical use of the here identified biomarker in assisting clinical diagnosis and treatment of ccRCC patients.

## Materials and methods

### Data and preprocessing

The TCGA transcript-expression level profiles (TPM and count values) of ccRCC and matched normal kidney samples was downloaded from https://osf.io/gqrz9 (Tatlow and Piccolo, 2016) on November 27, 2018, which was quantified by Kallisto (Bray et al., 2016) based on the GENCODE reference transcriptome (version 24). The clinical information of TCGA samples was downloaded through R package TCGAbiolinks (Colaprico et al., 2016). The whole-exome sequence data of 100 ccRCC samples of patients from Japanese cohort (Sato et al., 2013) was downloaded from European Genome-phenome Archive (accession number: EGAS00001000509). BEDTools (Quinlan and Hall, 2010) was used for converting BAM to FASTQ file. Kallisto was used for estimating the count and TPM values of transcripts based on the same reference transcriptome of TCGA data. The sum value of the multiple transcripts of a gene was used as the expression value of this gene. The genes with average TPM values >1 in ccRCC patients were analyzed.

### Differential expression analysis

DESeq2 (Love et al., 2014) was used to identify DEGs between two groups. The raw count values of genes were used as input of DESeq2. The Benjamini-Hochberg (BH) procedure was used to estimate FDR.

### Overlapping of two lists of DEGs

If DEG list 1 with *L*_*1*_ genes and DEG list 2 with *L*_*2*_ genes have *k* overlapping genes and *s* of these genes shows the same directions which means high expression of these genes indicates favorable/unfavorable survival or group 1/2 in both lists, the probability of observing at least *s* consistent genes by chance can be calculated according to the following cumulative hypergeometric distribution model:

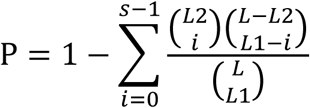

where *L* represents the number of the background genes commonly detected in the datasets from which the DEGs are extracted. The two DEG lists were considered to be significantly overlapping if P < 0.05.

The concordance score *s*/*k* is used to evaluate the consistency of DEGs between the two lists. Obviously, the score ranges from 0 to 1, and the higher concordance score suggests the better consistency of two lists of DEGs.

### Consensus clustering

Consensus clustering (Wilkerson and Hayes, 2010) was used for tumor classification based on the normalized expression profiles of signature genes by Z-score transformation. To achieve robust clusters, the data was resampled for 1000 times by considering 80% samples and signature genes resampling. The resampled data was transformed into a similarity matrix, termed as consensus matrix. K-means clustering was used to stratify samples based on the consensus matrix. The number of optimum cluster was determined by the lowest proportion of ambiguous clustering.

### Survival analysis

The univariate Cox regression model was used to evaluate the correlation of gene expression levels with OS. Survival curves were estimated by the Kaplan-Meier method and compared with the log-rank test.

### Functional enrichment analysis

GO enrichment was performed by the enrichGo function in R package ClusterProfiler (Yu et al., 2012), in which the hypergeometric distribution was used to calculate the statistical significance of biological pathways enriched with DEGs of interest.

### Development of the REOs-based biomarker

In each sample, the REO of every two signature genes (i and j) is denoted as either G_i_ > G_j_ or G_i_ < G_j_ exclusively, where G_i_ and G_j_ represent the expression values of gene i and j, respectively. For a given gene pair (G_i_ and G_j_), we used Fisher’s exact test to evaluate whether the frequency of group 1 samples with a specific REO pattern (G_i_ > G_j_ or G_i_ <G_j_) was significantly different from that in group 2 samples in each training dataset. The P values are adjusted using BH procedure. The gene pairs detected with 0.05 FDR control and over 70% difference of the frequency of their REOs between two groups were denoted as reversed gene pairs. The overlapped reversed gene pairs consistently identified from all the training datasets were selected as the candidate signature gene pairs. Totally, we found 171 signature gene pairs. For each signature gene pair, according to their within-sample REO, we classified the samples of each training dataset into high or low-risk groups and then evaluated the sensitivity and specificity of this gene pair. Here, the sensitivity is defined as the ratio of correctly identified high risk samples to all high-risk samples and the specificity is defined as the ratio of correctly identified low-risk samples to all low-risk samples. Then, from these signature gene pairs, we performed a forward selection procedure in each training dataset to search a set of gene pairs that achieved the highest F-score value, a harmonic mean of sensitivity and specificity, which is calculated as follows:

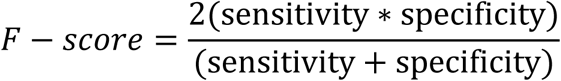

Taking each of the 171 gene pairs as a seed, we added another gene pair to the biomarker at a time until the F-score did not increase. The classification rule is that a sample is classified into high or low-risk group if the majority of the REOs of the set of gene pairs within this sample vote for high or low-risk. We got 171 biomarkers based on each training dataset. Totally, we got 17100 candidate biomarkers for all the 100 training datasets. Finally, we selected the biomarker with the lowest (1-F_1_)^2^+(1-F_2_)^2^+…+(1-F_n_)^2^ as the final biomarker, in which F_n_ is the F-score value in the n^th^ training dataset.

### Application of CMap2 data to drug discovery

The pre-procession of CMap2 data was described in our previous study (Turanli et al., 2019b). In brief, the gene expression profiles of three cell lines, HL60, MCF2 and PC3, were downloaded from https://portals.broadinstitute.org/cmap/ (CMap Build 02). For each cell line, gene Log2FC was used for comparison between treatment instances and its respective controls. Then, the confidence score was calculated per each drug-gene interaction using the P values from three cell lines. An approximation confidence score to 1 was assumed as the higher confidence level. The drug-gene pairs with CS>0.5 were used in further analysis.

## Supporting information

RNA catabolic process and nuclear transport pathways (FDR<0.05; Table S1).

## Acknowledgements

The study is funded by The Knut and Alice Wallenberg Foundation.

## Supplementary Table legend

Table S1. The enriched GO terms for the 539 signature genes

